# OrBITS: Label-free and time-lapse monitoring of patient derived organoids for advanced drug screening

**DOI:** 10.1101/2021.09.09.459656

**Authors:** Christophe Deben, Edgar Cardenas De La Hoz, Maxim Le Compte, Paul Van Schil, Jeroen M. Hendriks, Patrick Lauwers, Suresh Krishan Yogeswaran, Filip Lardon, Patrick Pauwels, Steven Van Laere, Annemie Bogaerts, Evelien Smits, Steve Vanlanduit, Abraham Lin

## Abstract

**Background:** Patient-derived organoids are invaluable for fundamental and translational cancer research and holds great promise for personalized medicine. However, the shortage of available analysis methods, which are often single-time point, severely impede the potential and routine use of organoids for basic research, clinical practise, and pharmaceutical and industrial applications.

**Methods:** Here, we developed a high-throughput compatible and automated live-cell image analysis software that allows for kinetic monitoring of organoids, named **Or**ganoid **B**rightfield **I**dentification-based **T**herapy **S**creening (OrBITS), by combining computer vision with a convolutional network machine learning approach. The OrBITS deep learning analysis approach was validated against current standard assays for kinetic imaging and automated analysis of organoids. A drug screen of standard-of-care lung and pancreatic cancer treatments was also performed with the OrBITS platform and compared to the gold standard, CellTiter-Glo 3D assay. Finally, the optimal parameters and drug response metrics were identified to improve patient stratification.

**Results:** OrBITS allowed for the detection and tracking of organoids in routine extracellular matrix domes, advanced Gri3D^®^-96 well plates, and high-throughput 384-well microplates, solely based on brightfield imaging. The obtained organoid Count, Mean Area, and Total Area had a strong correlation with the nuclear staining, Hoechst, following pairwise comparison over a broad range of sizes. By incorporating a fluorescent cell death marker, intra-well normalization for organoid death could be achieved, which was tested with a 10-point titration of cisplatin and validated against the current gold standard ATP-assay, CellTiter-Glo 3D. Using this approach with OrBITS, screening of chemotherapeutics and targeted therapies revealed further insight into the mechanistic action of the drugs, a feature not achievable with the CellTiter-Glo 3D assay. Finally, we advise the use of the growth rate-based normalised drug response metric to improve accuracy and consistency of organoid drug response quantification.

**Conclusions:** Our findings validate that OrBITS, as a scalable, automated live-cell image analysis software, would facilitate the use of patient-derived organoids for drug development and therapy screening. The developed wet-lab workflow and software also has broad application potential, from providing a launching point for further brightfield-based assay development to be used for fundamental research, to guiding clinical decisions for personalized medicine.

## 1. Introduction

The use of 2D cancer cell lines has traditionally been the gold standard in preclinical *in vitro* cancer research. However, these models fail to recreate the complex cell-cell interactions present in the tumor microenvironment and lack the genetic heterogeneity found in cancer patients. In addition, prolonged use of these cancer cell lines leads to acquired mutations or gene expression alterations, which are often overlooked as the cell line deviates from the originally derived tumor. As a result, new therapies are often met with high failure rates during translation from preclinical research to clinical trials, thus resulting in extreme forfeiture of research labor and financial loss.

The development of 3D cell culture technologies has greatly improved the physiological relevance of *in vitro* cancer models and the use of patient-derived organoids (PDOs) is revolutionizing basic and translation cancer research (1). These PDOs resemble both the pheno- and genotype of the tissue they are derived from and can be expanded long-term and cryopreserved to establish a living tumor and healthy tissue biobank (1). Furthermore, one of the most promising applications of PDOs is for personalized cancer treatment to predict clinical response *ex vivo* (2-8). However, the available repertoire of assays for high-throughput organoid analysis is severely limited, and therefore, the current gold standard relies on a rudimentary viability assay.

The gold standard analysis method, Promega CellTiter-Glo 3D cell viability assay, determines the number of viable cells in 3D cell cultures based on luminescent quantification of intracellular ATP in both 96- and 384-well microplate format. This method is met with several intrinsic limitations including inter-patient growth rate variations and (drug-induced) metabolic modulations, which could affect the translatability of this readout (9-11). For drug screening and diagnostic applications, this assay is also unable to determine the mechanistic action of the drug, such as cytostatic or cytotoxic response, and is limited to a single time-point analysis. These aspects critically limit the wide adoption of organoid technology for clinical diagnostics and pharmaceutical drug screening. Therefore, although the CellTiter-Glo viability assay has been a valuable method to monitor drug responses, more sophisticated high-throughput analysis methods are urgently needed.

In this study, we addressed these shortcomings by using an automated, high-throughput compatible, and kinetic screening platform to monitor therapy response. Using the Tecan Spark Cyto multi-mode plate reader system, we automated seeding of full-grown PDOs in a 384-well format and performed real-time, whole-well brightfield (BF) and fluorescence imaging in a temperature, CO_2,_ and O_2_ controlled environment. This reduces the variability involved in manual organoid cultures, which is highly cumbersome and not scalable. Using our in-house developed OrBITS (**Or**ganoid **B**rightfield **I**dentification-based **T**herapy **S**creening) live-cell image analysis software, which utilizes convolutional network machine learning to eliminate the use of fluorescent live-cell labelling dyes, we validated the capacity for BF imaging-based monitoring of PDO growth, by comparing our results to the current industry and research standards. The combination of BF imaging with a fluorescent cell death marker further demarcated cytostatic from cytotoxic therapy responses, thus providing greater insight into drug effects and the potential for improving translation into a durable clinical response in patients. Using this automation setup and OrBITS, we performed a screening of several chemotherapeutics and targeted therapies on patient-derived organoids as a proof-of-concept and identified the most suitable drug response metric. The technology described here to produce and validate our wet-lab workflow and image analysis software unlocks the potential for wide adoption of organoid-based assays for drug screening and discovery as well as guidance of clinical decision for personalized medicine.

## 2. Materials and Methods

### 2.1 Patient tissue

Tumor tissue and normal lung tissue (distant from the tumor site) were obtained from adeno- and squamous cell carcinoma non-small cell lung carcinoma (NSCLC) and pancreatic ductal adenocarcinoma patients undergoing curative surgery at the Antwerp University Hospital (UZA) in 2019-2021. Written informed consent was obtained from all patients, and the study was approved by the UZA Ethical Committee (ref. 17/30/339; 14/47/480). All samples were registered in the Biobank Antwerp, Belgium; ID: BE 71030031000. Mutation analysis on the parental NSCLC tissue was performed as part of standard-of-care using the Oncomine Focus Assay (Thermo Fisher Scientific) at the UZA pathology department.

### 2.2 Tissue processing and organoid culture

Tissue was stored in Ad-DF+++ (Advanced DMEM/F12 (GIBCO), with 1% GlutaMAX (GIBCO), 1% HEPES (GIBCO), 1% penicillin/streptomycin (GIBCO) supplemented with 2% Primocin (Invivogen) at 4°C and transported on ice to be processed within 24 hours for organoid culture. Tumor and normal tissue were minced with two scalpels, collected in 0.1% BSA precoated tubes and washed with PBS. Next, fragments were dissociated with 0.5 mg/mL dispase type II (Sigma-Aldrich), 1.5 – 5 mg/mL collagenase type II (Sigma-Aldrich), 1:500 Primocin and 10 µM Y-27632 (Cayman Chemicals) in MG^2+^/Ca^2+^ PBS (GIBCO) for 60 minutes at 37°C. Digested cells were washed three times with PBS and resuspended in 2/3 Cultrex Type 2 (R&D Systems) and 1/3 Full Ad-DF+++ medium and plated in drops which were allowed to solidify for 30 minutes at 37°C after which they were overlayed with Full Ad-DF+++ medium. Full lung Ad-DF+++ medium consisted of 10% Noggin conditioned medium (HEK293-mNoggin-Fc; kindly provided by Hans Clever, Hubrecht Institute), 10% R-spondin-1 conditioned medium (293T-HA-Rspol-Fc; kindly provided by Calvin Kuo, Stanford University), 1 x B27 supplement (GIBCO), 10 mM nicotinamide (Sigma-Aldrich), 25 ng/mM human recombinant FGF-7 (Peprotech), 100 ng/mL human recombinant FGF-10 (Peprotech), 500 nM A83-01 (Tocris), 1 µM SB202190 (Cayman Chemicals) and 5 µM Y-27632 (only used after passaging and thawing). Full PDAC Ad-DF+++ medium consisted of 0.5nM WNT Surrogate-Fc-Fusion protein (Immunoprecise), 4% Noggin-Fc Fusion Protein conditioned medium (Immunoprecise), 4% Rspo3-Fc Fusion Protein conditioned medium (Immunoprecise), 1x B27 without vitamin A, 10 mM nicotinamide, 100 ng/ml FGF-10, 500 nM A83-01, 10 nM gastrin (R&D Systems) and 5 mM Y-27632. For passaging, organoids were digested to single cells with TrypLE Express (GIBCO). For cryopreservation, 3-day old organoids were harvested with Cultrex Harvesting Solution (R&D Systems) and frozen in Recovery Cell Culture Freezing Medium (GIBCO). Samples were tested for Mycoplasma contamination with the MycoAlert Mycoplasma Detection Kit (LONZA).

### 2.3 Immunohistochemical analysis

Early passage organoids were collected using Cultrex Organoid Harvesting Solution (R&D systems), washed with ice-cold PBS, and fixated in 4% paraformaldehyde for 30 minutes at room temperature. Fixed organoids were transferred to a 4% agarose micro-array mold and paraffin-embedded as described before (12). Five µm-thick sections were prepared, deparaffinized and rehydrated prior to staining. Sections were subjected to heat-induced antigen retrieval by incubation in a low pH buffer (Envision Flex TRS low pH (DAKO) for 20 min at 97°C (PT-Link, DAKO). Endogenous peroxidase activity was quenched by incubation in peroxidase blocking buffer (DAKO) for 5 min. Slides were stained manually with mouse anti-TTF1 (clone SPT24; TTF-1-L-CE, Leica, 1/400, 25’) and mouse anti-P40 (clone BC28; Cat# ACI 3066 A, C, RRID:AB_2858274, Biocare, ready-to-use, 30’) primary antibodies followed by an incubation with ENVISION Flex+ Mouse Linker (DAKO, 15’) for signal amplification. Mouse anti-NapsinA (clone MRQ-60, Cell Marque, 1/350, 35’) staining was performed on a DAKO autostainer Link 48. After that the slides were incubated for 25 min with Envision FLEX/HRP (ready-to-use, DAKO) secondary antibody followed by 10 min incubation with the DAB substrate/chromogen detection system (DAKO). The sections were counterstained for 2 min with hematoxylin (0.1%), dehydrated and mounted with Quick-D Mounting Medium (KliniPath). Sections were imaged using a Leica DM500 microscope equipped with an ICC50 E camera.

### 2.4 In vitro drug screening

Three days before the start of the experiment, organoids were passaged as single cells using TrypLE and plated in Cultrex drops. Subsequently, organoids were harvested with Cultrex Harvesting Solution, collected in 15 mL tubes coated with 0.1% BSA/PBS, washed with Ad-DF+++ and resuspended in 1 mL Full Ad-DF+++ medium (without Y-27632). Next the number of organoids were counted with the Sceptor 2.0 using a 60 µM sensor (Merck Millipore). Organoids were then diluted in Full Ad-DF+++ and 5% Cultrex on ice to a concentration that results in 500-2000 organoids/60µL. 60µL of this solution was plated into a 384-well ultra-low attachment microplate (Corning, #4588) using the Tecan Spark Cyto Injector at a speed of 100 µL/s to avoid bubbles. All tubes and the Spark Cyto Injector were primed with 0.1 % BSA/PBS to avoid sticking of the organoids. Next, the plate was centrifuged (100 rcf, 30 sec, 4°C) and incubated for at least 30 minutes at 37°C.

All drugs and fluorescent reagents were added to the plate using the Tecan D300e Digital Dispenser. Cytotox Green (75 nM / well, Sartorius), Caspase 3/7 Green Reagent (2.5 µM / well, Sartorius), Z-VAD-FMK (50 µM / well, Bachem AG), Erlotinib, Gefitinib, Osimertinib and Afatinib (Selleckchem) were dissolved in DMSO. Hoechst 33342 (50 nM / well, ThermoFisher), Cisplatin (Tocris) and Carboplatin (Selleckchem) were dissolved in PBS to yield a final concentration of 0.3 Tween-20 required for dispensing with the D300e Dispenser.

BF, green and blue fluorescence whole-well images (4x objective) were taken with the Tecan Spark Cyto set at 37°C / 5% CO_2_ for kinetic experiments in a humidity cassette. For endpoint measurement of ATP levels, 60 µL CellTiter-Glo 3D reagent (Promega) was injected using the Tecan Spark Cyto Injector to each well, shaken for 5 minutes and measured after 30 minutes incubation with the Tecan Spark Cyto luminescence module.

A detailed video protocol will be made available in the Journal of Visualized Experiments (13).

### 2.5 Drug response metrics

Dose-response curves were plotted and IC_50_-values were calculated using GraphPad Prism 9 (GraphPad Prism, RRID:SCR_002798). Drug concentrations were transformed to log10 and raw data results were normalized to vehicle (100%) and/or baseline control (0%) (Staurosporin 5 µM or 100 µM cisplatin) for viability assessment, and vice versa for cell death assessment (EC_50_). Curves were fitted using the log (inhibitor/agonist) vs. normalized response - Variable slope function.

The growth rate (GR) metric (negative control) and the normalized drug response (NDR) metric (positive and negative controls) were calculated based on the work of Hafner and colleagues (14) and Gupta and colleagues (15), respectively, using the adapted R script from Gupta et al.. The Total BF Area – Total Green Area parameter was used, and the fold change was calculated for each well individually from the first measurement (T0) and a timepoint as indicated in the figures. Based on the NDR values, the drug effects can be classified as: >1, proliferative effect; = 1, normal growth as in negative control; = 0, complete growth inhibition; = −1, complete killing. Screen quality was determined by calculating the Z Factor score using the formula

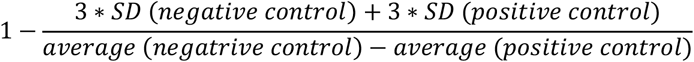

### 2.6 Image-based analysis

Following image acquisition with the Tecan Spark Cyto, BF and fluorescence images were analyzed using an in-house developed analysis software OrBITS. Briefly, a fully convolutional network was used to take single images and return sematic segmentation maps as outputs. The model training methodology was performed by using fluorescence intensity images as ground truth. A detailed description about the image analysis and training methodology is provided in Supplementary Information. An image mask, a colored overlay of identified objects, is applied to visualize the accuracy of the image segmentation. The analysis output from OrBITS (Table S1) was correlated with the current gold standard assays: Hoechst staining (for organoid tracking), Cytotox Green (for organoid death), and the CellTiter-Glo 3D assay (for drug screening). Use of the OrBITS image analysis software we developed is available to researchers upon reasonable request and discussion with the corresponding author. In addition, a web-based platform is currently being developed to increase accessibility of the software to users. Interest in this platform can be expressed to the corresponding author.

### 2.7 Organoid mutation analysis

Full grown organoids were collected as described above and DNA was isolated using the QIAamp DNA blood mini kit (Qiagen) and send to Genewiz Europe (Leipzig, Germany) for whole exome sequencing on an Illumina NovaSeq platform (2 × 150 bp sequencing, 12Gb (120x)). Raw reads were mapped onto the human reference genome (hg38) using the *BWA MEM* algorithm (v.0.7.17) in standard settings. Resulting SAM-files were converted into BAM-files, coordinated-sorted, and indexed using *samtools* (v.1.9). Reads overlapping with 86 genes mutated in at least 10% of either PDAC or NSCLC samples included in the cBioPortal for cancer genomics were selected using the *bedtools* (v.2.29.2) intersect-command retaining the full read in the original BAM-file (-wa option). Variants were called using the haplotype-based variant detector freebayes (v.1.3.2; Garrison E, Marth G. Haplotype-based variant detection from short-read sequencing. arXiv preprint arXiv:1207.3907 [q-bio.GN] 2012) in standard settings and indel positions were left-aligned and normalized using the *bcftools* (v.1.9) norm-command. Resulting VCF-files were annotated using *SnpEff* (v.5.1) for functional annotations and *SnpSift* (v.5.1) for human genetic variation using dbSNP (build 154), ClinVar (release 05/09/2022) and COSMIC (v96). Annotated VCF files were further manipulated using the *SnpSift* filter-command retaining only coding, non-synonymous variants that are contained COSMIC, that are not considered as benign and/or likely benign by ClinVar, that have a minimal coverage of 120x with at least 40 reads supporting the alternate allele with no positional or strand bias, that are not of germline origin or are reported as common variants (minor allele frequency>1%) in any human population and that are either missense, stop gained or frame shift mutations. Mutational profiles were visualized in oncoprint format.

### 2.8 Statistical Analysis

Correlations were measured and visualized using linear fit regression using the least squares approximation. The equation reports slope and bias between the two measured values as well as the R squared value. Log IC50-values were compared using the extra sum-of-squares F test in GraphPad Prism with a significance cut-off of p < 0.05.

## 3. Results

### 3.1 Establishment of patient-derived organoids

Validation of the organoid monitoring ability using BF imaging was performed on various lung organoid lines. Organoids derived from NSCLC patient tumor resection fragments had a predominantly solid growth pattern, while organoids derived from distant healthy lung tissue displayed a cystic growth pattern (Fig. S1). Further immunohistochemical characterization of the tumor tissue-derived organoids indicated that the cells were non-malignant and most likely represent metaplastic squamous epithelial cells as suggested by Dijkstra and colleagues (details in supplementary information, Fig. S1 and Table 1) (17, 18). This was further confirmed genetically, since NSCLC_006T and NSCLC_013T organoids lack the KRAS and EGFR mutation found in the parental tumor, respectively (Fig. S2, Table 1). These observations confirmed the challenges related to generating pure NSCLC organoids. However, for validation of organoid monitoring with OrBITS, these non-malignant organoids were equally relevant. The cystic NSCLC_006N and NSCLC_051N organoids and solid NSCLC_013T, NSCLC_046T and NSCLC_051T organoids were used in the subsequent experiments. In addition, the KRAS mutant PDAC_052 and PDAC_060 lines were included as genetically validated tumor organoids (Fig. S2, Table 1).

**Table 1:** Overview of patient-derived organoid lines

| Table 1: Overview of patient-derived organoid lines |  |  |  |  |  |  |  |  |
| --- | --- | --- | --- | --- | --- | --- | --- | --- |
| Sample | Source | Morphology | Napsin A | P40 | TTF1 | Mutation<br>Parental<br>Tumor | Mutation in<br>Organoids | Organoids |
| NSCLC_006N | Normal | Cystic | -/- | +/-<br>(polarized) | +/+ | NA | NA | Normal |
| NSCLC_006T | AC | Solid/Cystic | -/- | +/-<br>(polarized) | +/+ | KRAS<br>c.34G<T | No | Non-malignant |
| NSCLC_013N | Normal | Cystic | +/- | +/-<br>(polarized) | +/+ | NA | NA | Normal |
| NSCLC_013T | AC | Solid | -/- | +/-<br>(polarized) | +/+ | EGFR<br>c.2573T>G | No | Non-malignant |
| NSCLC_046N | Normal | Cystic | -/- | +/-<br>(polarized) | +/+ | NA | NA | Normal |
| NSCLC_046T | AC | Solid | -/- | +/+ | +/+ | No | No | Non-malignant |
| NSCLC_051N | Normal | Cystic | -/- | +/-<br>(polarized) | +/+ | NA | NA | Normal |
| NSCLC_051T | SQ | Solid | -/- | +/-<br>(polarized) | +/+ | No | No | Non-malignant |
| PDAC_052 | AC | Cystic | NA | NA | NA | NA | KRAS<br>c.34G>T | Tumoral |
| PDAC_060 | AC | Cystic | NA | NA | NA | NA | KRAS<br>c.35G>A | Tumoral |
| NSCLC: non-small cell lung cancer; PDAC: pancreatic ductal adenocarcinoma; AC: adenocarcinoma; SQ: squamous cell carcinoma; NA: not applicable. |  |  |  |  |  |  |  |  |

### 3.2 Validation of label-free Brightfield monitoring of all stages of organoid growth

A kinetic image set was generated from a serial dilution of single cells and 3-day old organoids grown in a 384-well microplate from two organoid lines (NSCLC_013T and NSCLC_006N) stained with Hoechst as a gold standard reference method. Pairwise comparison showed a strong correlation of the organoid Count, Mean Area, and Total Area detected by BF when compared to Hoechst for the entire range of organoid sizes (Fig. 1A). This indicates that OrBITS is capable of kinetic monitoring at all stages of organoid growth in the absence of a fluorescent dye and based solely on BF image analysis (Fig. 1B, Supplemental Video 1-3). For Mean and Total Mask Area, BF analysis resulted in a higher area compared to Hoechst analysis, mainly in the cystic NSCLC_006N line since Hoechst only identifies the edge of cystic organoids (Fig. 1A-B).

**Figure 1:**
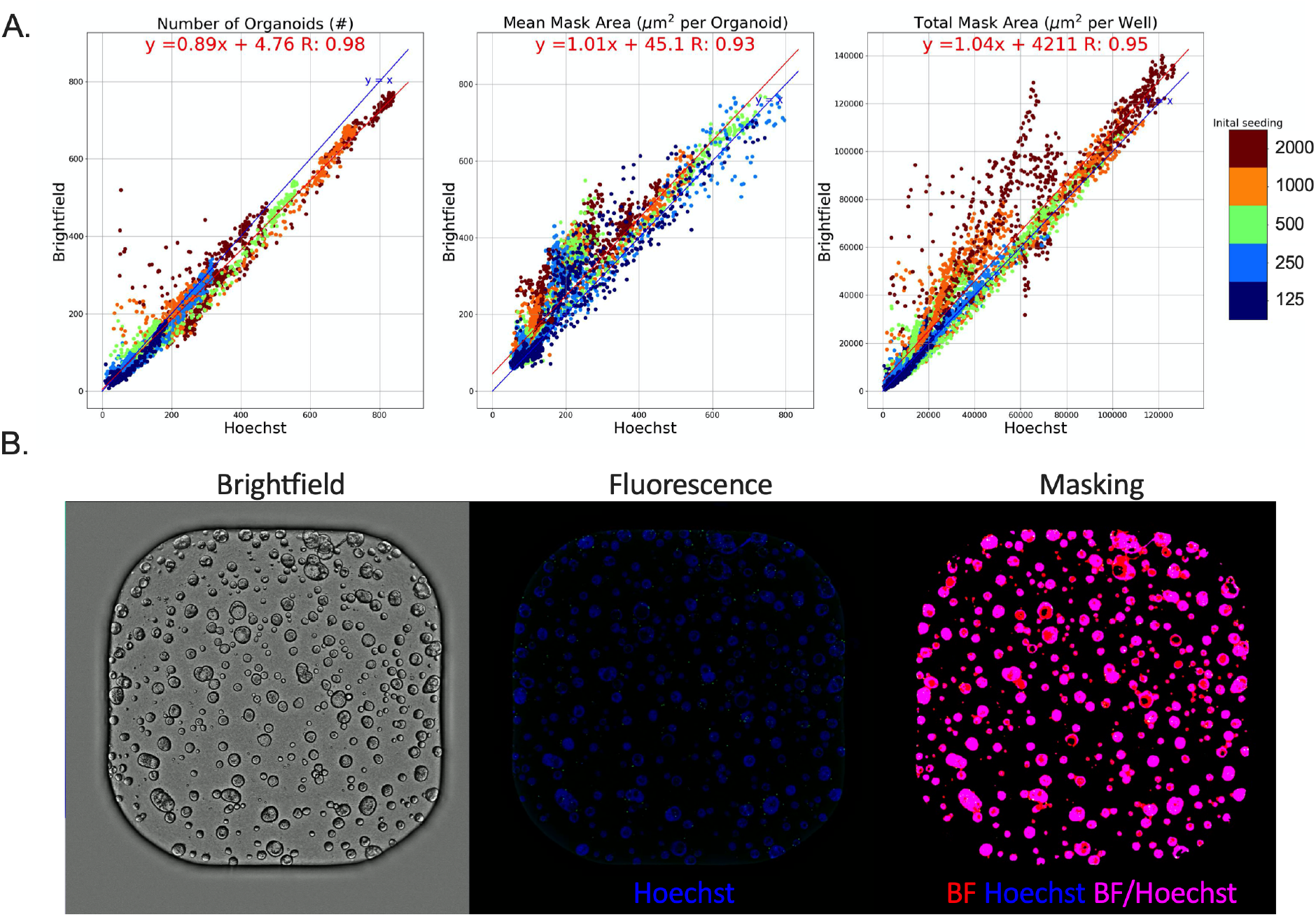
Brightfield versus Hoechst. (A) Pairwise comparison of organoid Counts, Mean Mask Area and Total Mask Area detected by Brightfield- and Hoechst-based analysis of NSCLC_013T and NSCLC_006N PDO lines plated as a serial dilution of single cells and 3-day old organoids (Table S2). Blue line: line of unity (x=y), red line: best-fit linear regression. (B) Representative whole 384-well brightfield and fluorescent images of full-grown PDOs (NSCLC_006N) and the corresponding analysis mask. Supplemental Video 3 presents the corresponding time-laps video.

Both organoid lines could be grown from either single cells (Fig. 2A) or fully-grown organoids, which are more conventionally used (Fig. 2B). An important advantage of the proposed assay is that growth kinetics can be accurately monitored using only a limited amount of starting material. In fact, using a larger number of single cells or organoids affected BF image monitoring results in two ways: (i) when both lines (NSCLC_013T and NSCLC_006N) were plated at high single cell densities, a drop in counts was observed over time due to organoids merging or fusion events (Fig. 2C-D, Supplemental Video 1) resulting in an aberrant growth rate based on Mean Area for NSCLC_013T at higher seeding densities (Fig. 2A); (ii) NSCLC_006N PDOs showed a higher growth rate compared to NSCLC_013T PDOs due to differential nutrient requirements. At higher concentrations (>500 counts), we clearly observed a drop in NSCLC_006N organoids growth rate compared to the slower growing NSCLC_013T organoids (Fig. 2B).

**Figure 2:**
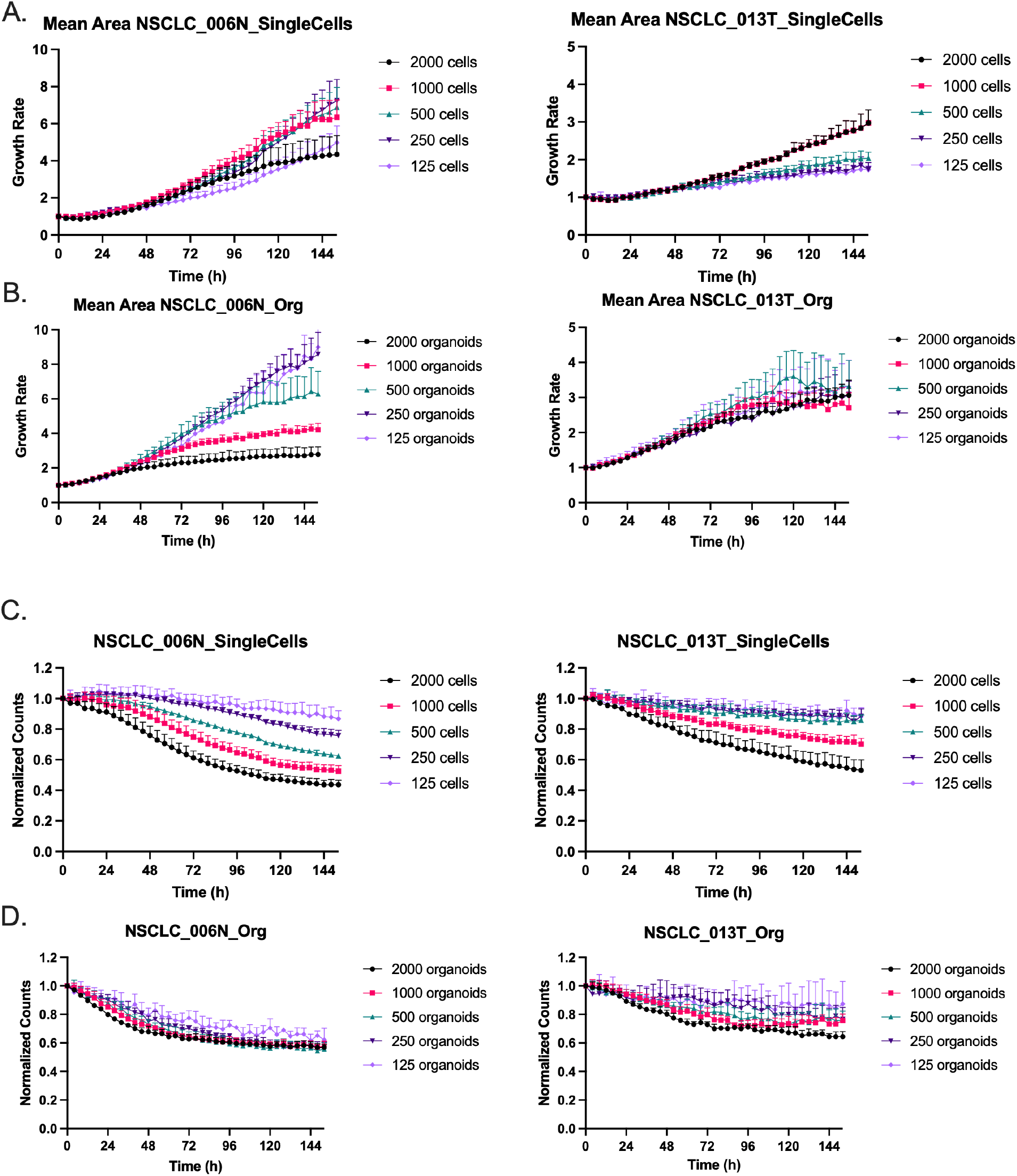
Influence of seeding density. Average growth rate of PDOs seeded as (A) single cells or (B) 3-day old organoids. Normalized counts (h=0) of PDOs seeded as (C) single cells or (D) 3-day old organoids. A range between 125 – 2000 cells or organoids were plated. Graphs presented as mean ± SD of 5 technical replicates and images were acquired every 6 hours.

### 3.3 Monitoring organoid death with intra-well normalization

Since PDOs used in drug screening will undergo varying degrees and stages of death, it is critical that the OrBITS software is able to detect such organoids. Therefore, a second kinetic image set was generated from two additional organoid lines to increase variation (NSCLC_051T and NSCLC_051N; 500/well plated as 2-day old organoids) and treated with a 10-point titration of cisplatin. PDOs were stained with Hoechst as the gold standard reference method and used for validation of BF image analysis. Despite varying levels of cytotoxicity from the chemotherapeutic treatment, the data showed that organoids were successfully identified from BF images (Fig. S3). Importantly, common artefacts (e.g. air-bubbles, extracellular matrix, dust) did not disrupt BF analysis (Fig. S3).

To further develop the applicability of the OrBITS software, we investigated the capacity for intra-well normalization of organoid death with cell death markers, such as Cytotox Green. Since Hoechst and Cytotox Green are both nuclear stains, the overlap of Green Area or Green Intensity (relative fluorescence units; RFU) with Hoechst allowed for intra-well normalization to study varying levels of cell death from the chemotherapeutic treatment (Fig. 3A). Importantly, pairwise comparison showed a strong correlation between BF and Hoechst normalized Green Area and Green Intensity, thus making a fluorescent live-cell labelling dye (nuclear or cytoplasmatic) redundant (Fig. 3B). Total Green Area / Total Mask Area resulted in the broadest dynamic range (∼Δ0.6 vs. ∼Δ0.4), although a complete overlap (R = 1) in the 100 % cell death control (100 µM CDDP) was not reached (Fig. 3B). Therefore, the proposed analysis provides an important advantage by including intra-well, normalized organoid death, even without the use of a viability label. Overall, we showed that our OrBITS software can mask organoids with high accuracy ranging from single cells to full-grown organoids based on BF imaging. In addition, inclusion of a fluorescent cell death marker allowed for kinetic detection of intra-well normalized organoid death. This allows for the removal of a nuclear or cytoplasmatic viability stain during kinetic analysis, a key limitation of the current state-of-the-art analysis, as it can influence cellular growth and health and confounds the effect of treatments.

**Figure 3:**
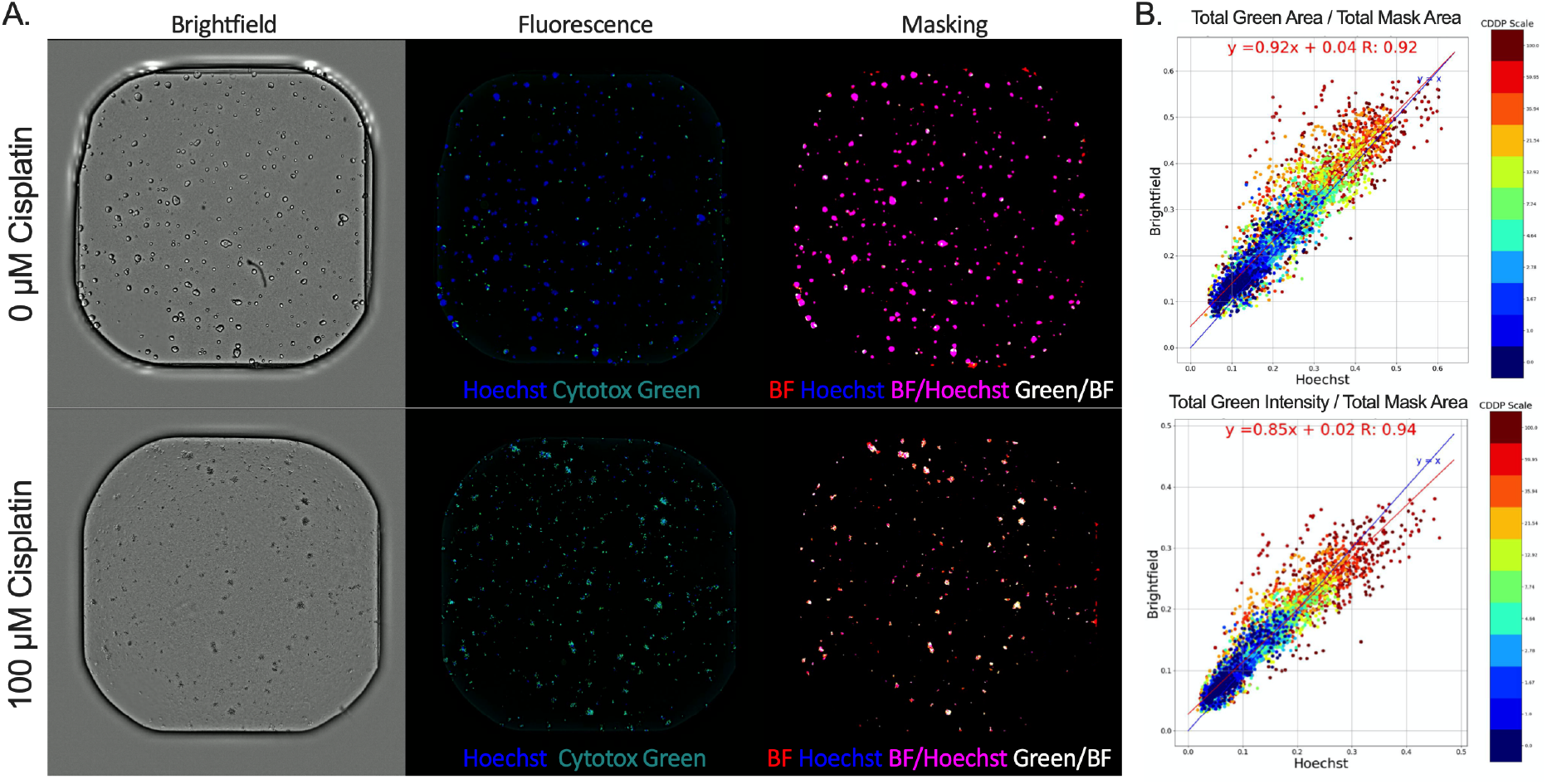
Brightfield vs Hoechst normalized fluorescent cell death marker. (A) Representative brightfield and fluorescent (Hoechst and Cytotox Green) images of untreated and 100 µM cisplatin (CDDP) treated NSCLC_051T PDOs and the corresponding masking of Brightfield, Hoechst, Brightfield/Hoechst overlap and Cytotox Green/Brightfield overlap. (B) Pairwise comparison of Total Green Area / Total Mask Area (per well: µm^2^/ µm^2^) and Total Green Intensity / Total Mask Area (per well: RFU/ µm^2^) with Mask Area based on either Hoechst fluorescence imaging or BF imaging (Table S2). Blue line: line of unity (x=y), red line: best-fit linear regression.

### 3.4 Validation of OrBITS brightfield PDO monitoring in proof-of-concept drug screening: Comparison to CellTiter-Glo3D

In order to determine if the OrBITS software could accurately monitor viability and therapy response in PDOs, we compared the BF image analysis with the current industry and research standard: CellTiter-Glo 3D viability assay. PDOs were again treated with a 10-point titration of cisplatin, imaged and analyzed with both OrBITS and the CellTiter-Glo 3D viability assay, and the read-outs were compared. Pairwise comparison revealed that the Total Mask Area had the strongest correlation with the CellTiter-Glo 3D luminescent signal, since both parameters result from whole-well readouts (Fig. 4A).

**Figure 4:**
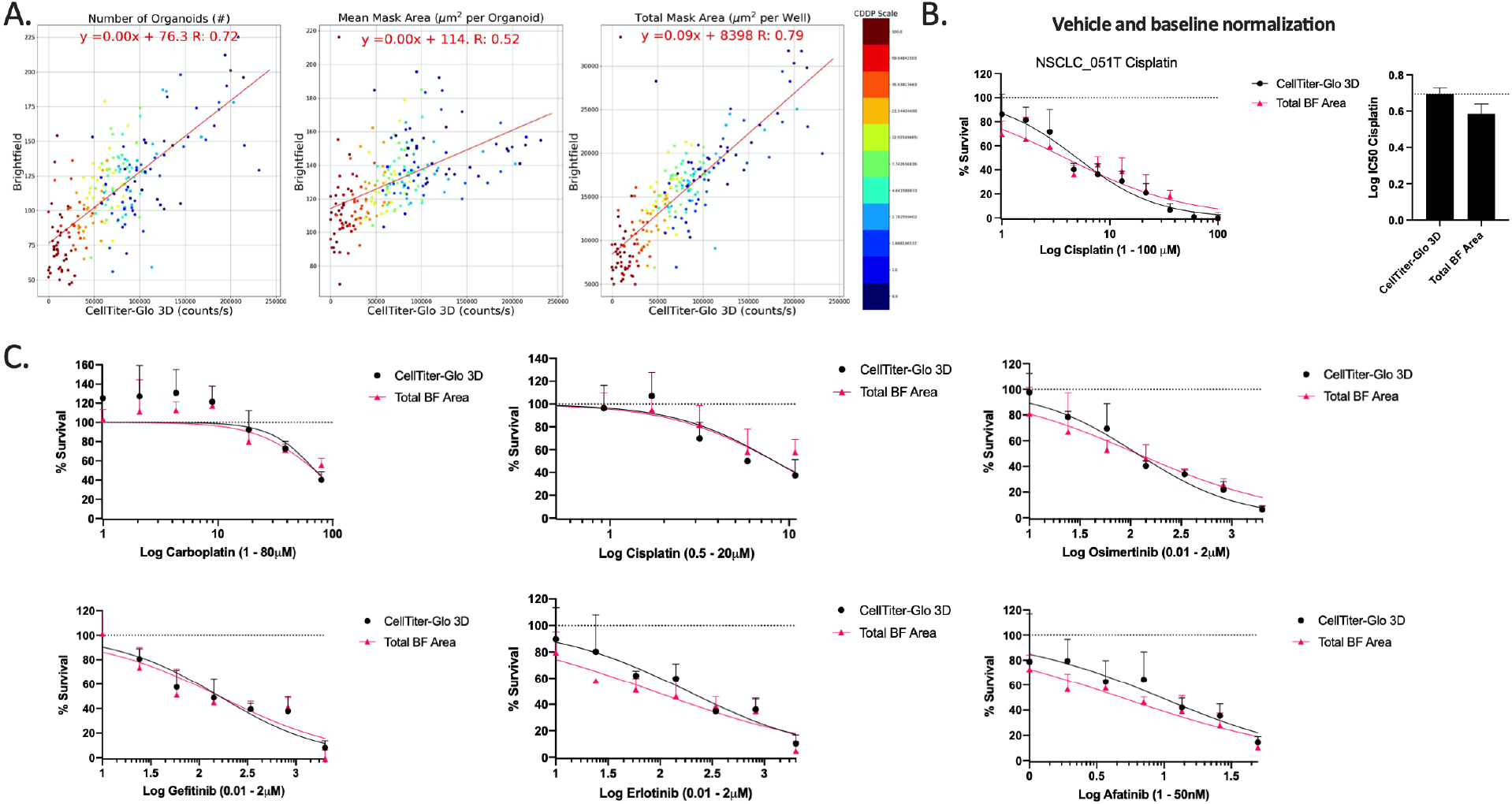
Brightfield imaging as a viability marker to monitor therapy response. (A) Pairwise comparison of organoid Count, Mean Mask Area and Total Mask Area with the luminescent read-out (counts/s) of the CellTiter-Glo 3D assay for the NSCLC_051T, NSCLC_051N, co-culture NSCLC_051T/N and NSCLC_050N PDO lines treated with a 10-point titration of cisplatin. Both read-outs were acquired at the same timepoint (114 hours) (Table S2). (B) Dose-response curve and corresponding IC50-values for NSCLC_051T treated with a 10-point titration of cisplatin and normalized to vehicle (100%) and baseline (0%, 100 µM cisplatin). (C) Dose-response curves of NSCLC_013T PDOs treated with a 7-point titration of cisplatin, carboplatin, erlotinib, gefitinib, osimertinib or afatinib normalized to vehicle (100%) and baseline (0%, 1 µM staurosporine). Corresponding IC50 values are presented in fig. S4. Graphs presented as mean ± SD of 5 technical replicates.

The assay quality was assessed based on the Z-factor, a coefficient reflective of both assay signal dynamic range and data variation, using 100 µM cisplatin as positive control for NSCLC_051T. The CellTiter-Glo 3D read-out was characterized by a Z-factor of 0.78 and the Total BF Area by a Z-factor of 0.69, indicating that the BF assay is excellent for drug screening. When applying both vehicle (100% viability; PBS) and baseline normalization (0% viability; 100 µM cisplatin) no significant difference (p = 0.0791) was observed between IC50 values obtained by the Total BF Area analysis and CellTiter-Glo 3D assay (Fig. 4B).

To validate our approach, we performed a drug screen of 6 standard-of-care lung cancer treatments on an additional PDO line (NSCLC_013T) using 1 µM staurosporine as the standardized 100% cell death control. A clear overlap of the inhibitory dose-response curves (Fig. 4C) and corresponding IC50 values (Fig. S4) was observed between the Total Mask Area analysis and CellTiter-Glo 3D assay. Therefore, it was clear that the OrBITS software could accurately monitor PDOs for various anti-cancer drug screenings.

We observed that the Total BF Area resulted in an overestimation of the percentage of viable cells compared to the CellTiter-Glo 3D assay, when limiting normalization to the vehicle control (Fig. 5A-C). The difference increased with cisplatin concentration, which was not unexpected since dead organoids were still counted by BF imaging. Therefore, we used a fluorescent cell death marker (Cytotox Green or equivalent) to correct for the area covered by dead entities, thus producing the parameter Total BF Area – Total Green Area. This parameter was characterized by a Z-factor of 0.80 for the NSCLC_051T line and both the dose-response curves and IC50-values showed an improved overlap with the CellTiter-Glo 3D assay for NSCLC_051T, 051N/T (mixed N and T) and 051N organoids (Fig. 5A-C). Importantly, the parameter showed a clear distinction in sensitivity between the NSCLC_051T and NSCLC_051N lines, with an intermediate response in the co-culture of both lines (Fig. 5D). In addition, this parameter allowed for kinetic monitoring of cell viability (Fig. 5E) and vehicle-normalized survival (Fig. 5F), thus making early detection of therapy response possible.

**Figure 5:**
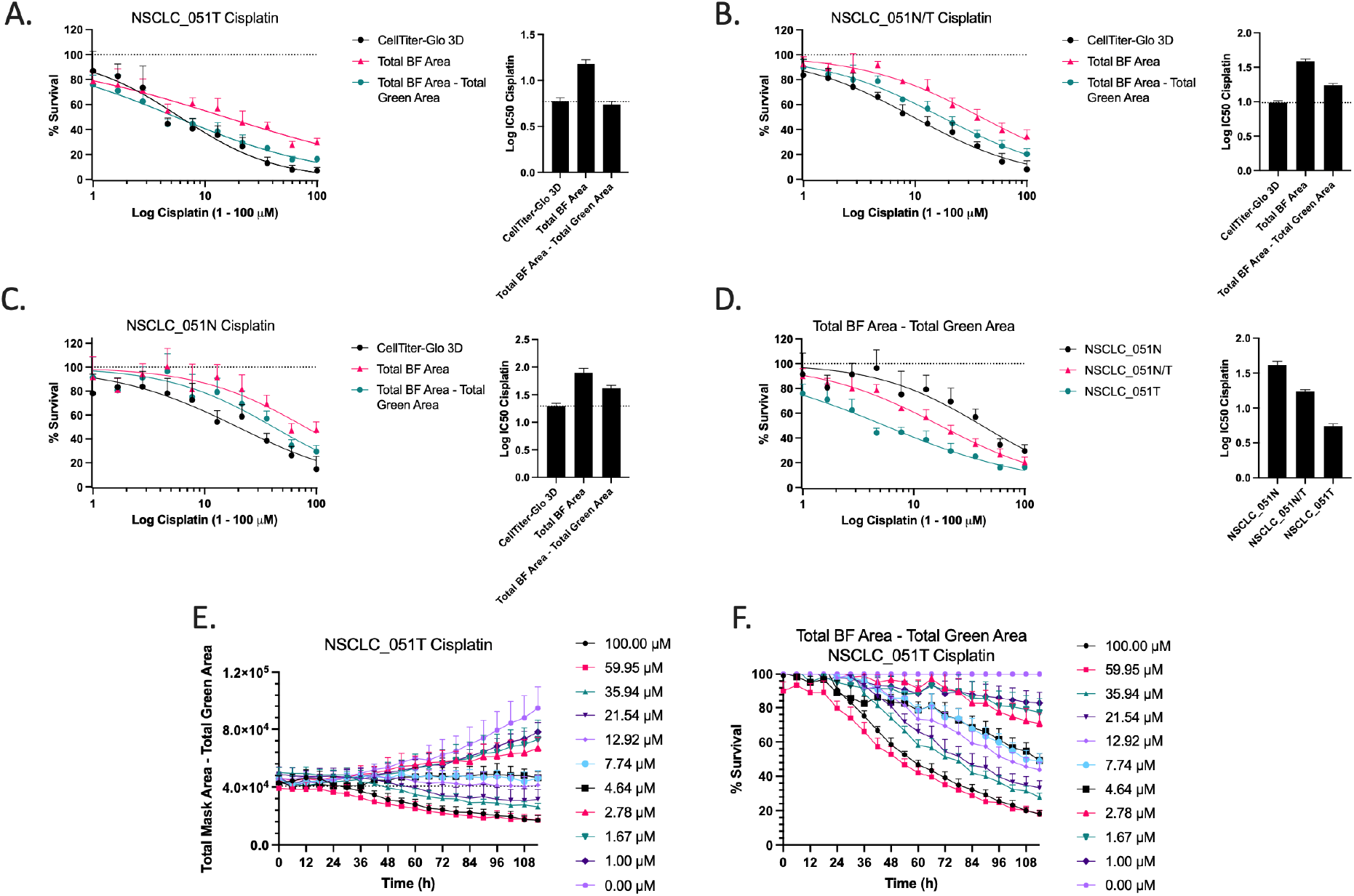
Cell death corrected vehicle normalization. (A-C) Dose-response curves and corresponding IC50-values for NSCLC_051T, NSCLC_051N and co-culture NSCLC_051N/T treated with a 10-point titration of cisplatin and normalized to vehicle (100%) control. Real-time monitoring of (E) cell viability and (F) survival percentage of NSCLC_051T PDOs treated with a 10-point titration of cisplatin. Percent survival was baseline corrected to vehicle control at each timepoint. Graphs presented as mean ± SD of 5 technical replicates.

Overall, we demonstrated that the parameters used in our proposed BF image analysis were robust and determined cell viability and IC50-values corresponding to the gold standard CellTiter-Glo 3D assay. Our OrBITS software is a significant advance from the current gold standard assay by addressing its intrinsic limitations, as it allows for kinetic monitoring of organoid health and growth from limited starting material and independent of metabolic changes.

### 3.5 Validation of OrBITS analysis to delineate cytostatic and cytotoxic anti-cancer therapy response

The ATP-based PDO drug screening analysis method does not provide insight into the mechanistic action of the drug, and therefore, we aimed to distinguish a cytostatic response (i.e. growth arrest) from a cytotoxic response (i.e. cell death) using the OrBITS software. To achieve this, we quantified the amount of therapy-induced organoid death using our BF and fluorescent image analysis and the CellTiter-Glo 3D assay. Pairwise comparison showed an inverse correlation of Total Green Area and Total Green Intensity normalized to Total BF Area with the CellTiter-Glo 3D readout (Fig. 6A). The assay quality was assessed based on the Z-factor using 100 µM cisplatin as positive control for NSCLC_051T.Total Green Area / Total BF Area and Total Green Intensity / Total BF Area were characterized by a Z-factor of 0.71 and 0.75, respectively, consistent with a robust assay. A clear inverse relation was observed for the inhibitory dose-response curve (% survival) and stimulatory dose-response curve of both the Total Green Area and Intensity parameters normalized to BF area (% cell death) for cisplatin treated NSCLC_051T PDOs, suggesting a cytotoxic response in this organoid line (Fig. 6B). An example of a cytostatic response is given for NSCLC_013T treated with gefitinib (Fig. 6C). The use of Total Green Area was a more standardized parameter since it was less susceptible to variability compared to the Total Green Intensity parameter (e.g. led intensity, exposure time, and dye concentration). In addition to delineating cytotoxic and cytostatic therapy responses, our method allowed for kinetic monitoring of therapy-induced cell death (Fig. 6D) and assessment of organoid health (growth and viability) as an imperative run quality control (Fig. 6E). Moreover, our method allowed for more in-depth research on the type of therapy-induced cell death that occurred. For example, cisplatin induced caspase 3/7 activity in NSCLC_046T, which is partly inhibited by the pan-caspase inhibitor Z-VAD-FMK (Fig. 6F). Consistently, Z-VAD-FMK inhibited cisplatin induced cell death, indicating that cytotoxicity occurs through caspase-dependent apoptosis (Fig. 6G). Here, the importance of kinetic monitoring is further highlighted, as the inhibitory effect of Z-VAD-FMK is nullified at 72h.

**Figure 6:**
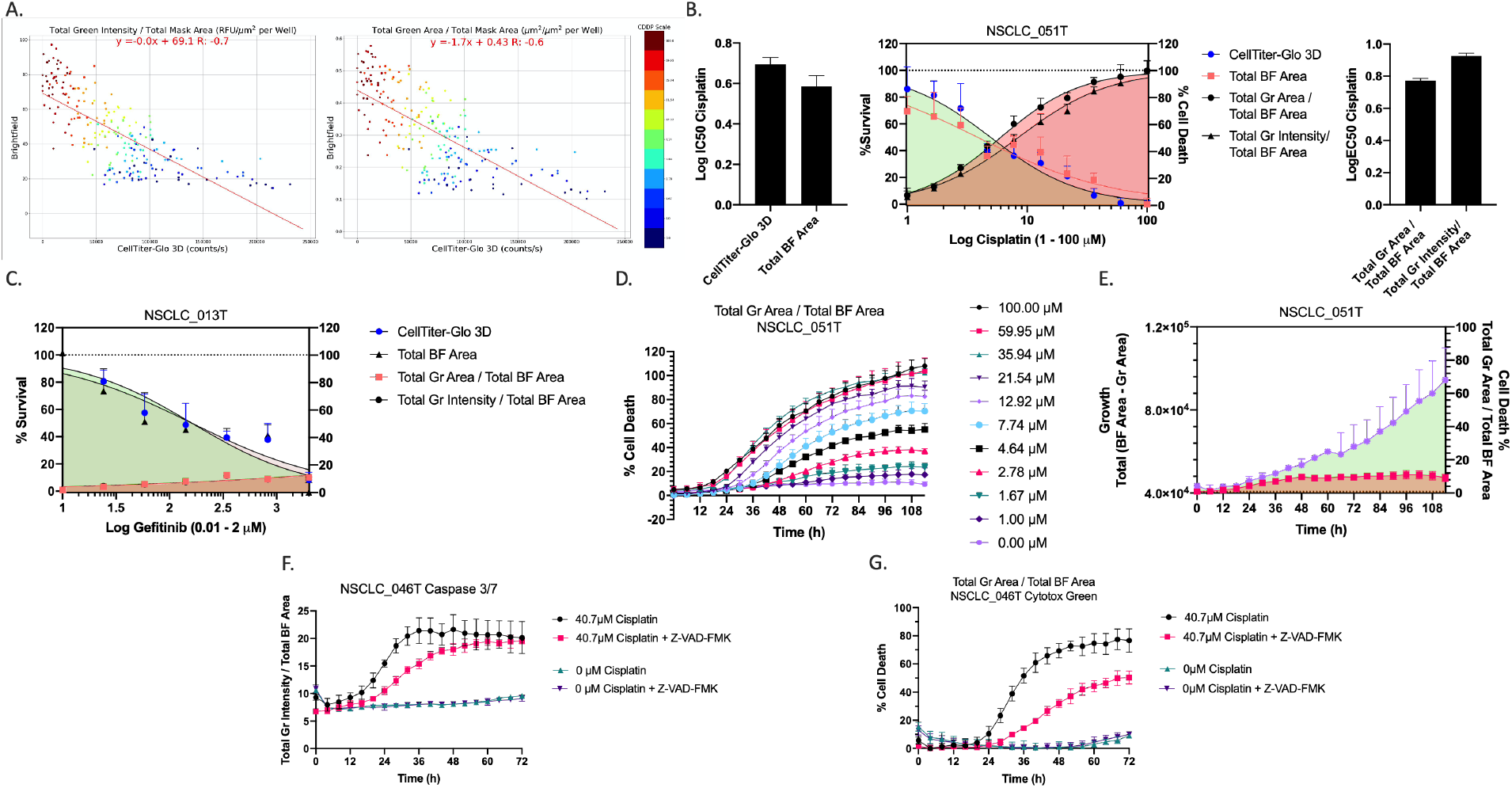
Fluorescent/brightfield imaging as a cell death marker. (A) Pairwise comparison of Total Green Intensity and Total Green Area (Cytotox Green reagent) normalized to Total BF Area with the luminescent read-out (counts/s) of the CellTiter-Glo 3D assay for the NSCLC_051T, NSCLC_051N, co-culture NSCLC_051T/N and NSCLC_050N PDO lines treated with a 10-point titration of cisplatin (Table S2). (B) Inhibitory (% survival, normalized to vehicle (100%) and baseline control (0%, 100 µM cisplatin) and stimulatory (% cell death, normalized to vehicle (0%) and positive control (100%, 100 µM cisplatin) dose-response curves and the corresponding IC50 and EC50 values for NSCLC_051T PDOs treated with a 10-point titration of cisplatin. (C) Inhibitory (% survival, normalized to vehicle (100%) and baseline control (0%, 1 µM staurosporine) and stimulatory (% cell death, normalized to vehicle (0%) and positive control (100%, 1 µM staurosporine) dose-response curves NSCLC_013T PDOs treated with a 7-point titration of gefitinib. (D) Real-time monitoring of therapy-induced cell death of NSCLC_051T PDOs treated with a 10-point titration of cisplatin. Total Green Area / Total BF Area was normalized to baseline (0%, vehicle at t = 0h) and positive (100%, 100 µM cisplatin at t = 72h) controls. (E) Real-time monitoring of organoid growth (Total BF Area – Total Green Area) and cell death (Total Green Area / Total BF Area) as run quality control. (F) Total Green Intensity / Total BF Area of the Caspase 3/7 green reagent in NSCLC_046T PDOs treated with cisplatin +/-Z-VAD-FMK (pan-caspase inhibitor). (G) Percentage of cell death of NSCLC_046T organoids treated with cisplatin +/-Z-VAD-FMK. Total Green Area / Total BF Area was normalized to baseline (0%, lowest value) and positive (100%, 75 µM cisplatin at t = 72h) controls. Graphs presented as mean ± SD of 5 technical replicates.

Overall, we demonstrated that OrBITS allowed for kinetic and endpoint analysis of therapy-induced cell death to distinguish cytostatic from cytotoxic responses and can be further developed to provide in-depth insight into the mechanistic action.

### 3.6 The normalized drug response metric improves patient stratification

Inter-patient differences in PDO growth rate and experimental artifacts like variations in inter-well seeding densities can lead to biased drug response estimates. Growth rate-based metrics correct for these variations and our method allows for the quantification of intra-well growth rate over time. When comparing the GR (negative control, fig. 7A) and NDR (positive and negative control, fig. 7B) metrics to the cell death percentage of cisplatin-treated NSCLC_051T PDOs, we identified that the NDR calculated from Total BF Area – Total Green Area most accurately resembles the cytotoxic response. In addition, the NDR (Fig. 7D) improved the separation between gemcitabine drug response from two PDAC PDOs compared to GR (Fig. 7C), which more accurately represented the clinical response of the patients treated with neoadjuvant gemcitabine-paclitaxel since PDAC_052 had a prolonged clinical response compared to PDAC_060 (disease control of 1 year vs. 4 months). The potential value of our method and the use of the NDR metric for improved therapy response prediction is currently being validated in a larger cohort of patients.

**Figure 7:**
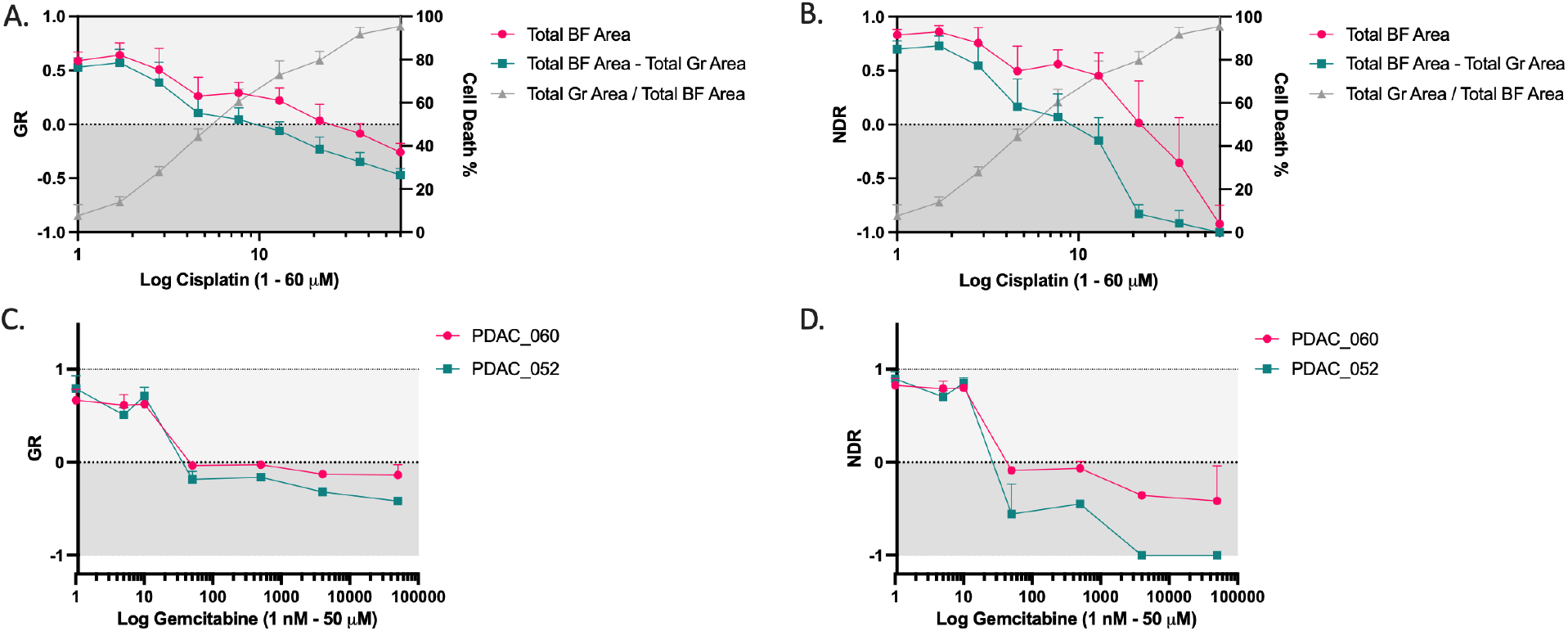
Growth Rate (GR) and normalized drug response (NDR) metrics. (A) NSCLC_051T treated with a 9-point titration of cisplatin. Left y-axis: GR (t = 120h) normalized to vehicle control. Right y-axis: % cell death, normalized to vehicle (0%) and positive (100%, 100 µM cisplatin) control. (B) NSCLC_051T treated with a 9-point titration of cisplatin. Left y-axis: NDR (t = 120h) normalized to vehicle and positive (100 µM cisplatin) control. Right y-axis: % cell death, normalized to vehicle (0%) and positive control (100%, 100 µM cisplatin). PDAC_060 and PDAC_052 treated with a 7-point titration of gemcitabine and (C) GR and (D) NDR calculated from Total BF Area – Total Green Area and normalized to vehicle control and vehicle and positive control (2 µM staurosporine), respectively.

### 3.7 Broad applicability of OrBITS for various organoid platforms and cancer cell lines

While the OrBITS software was developed using lung organoids in a high-throughput 384-well plate format, we have also successfully implemented the image analysis method for a range of cancer cell lines and PDOs of other tumor types. For example, we demonstrated the precise tracking capability on the growth of 3 cell lines (NCI-H1975, HCT-15, and NCI-H460) and pancreatic cancer-derived organoids using BF imaging (Supplemental Video 4). An overview off compatible cell lines we have tested to date is provided in table S3. Furthermore, we demonstrated that OrBITS is able to accurately mask organoids in other platforms. Pancreatic cancer organoids embedded in extracellular matrix domes were successfully monitored with OrBITS (Fig. 8A, Supplemental Video 5). This is highly advantages as it allows for monitoring of routine organoid cultures and long-term screening assays in various culture plate. OrBITS BF image analysis was also tested on the Gri3D^®^-96 well plates, an organoid plate format with hydrogel-based microcavities for high-throughput organoid cultures(19). Impressively, OrBITS accurately discriminated organoids from background in the BF images without additional training of the convolutional neural network (Fig. 8B, Supplemental Video 6). Taken together, it is clear that OrBITS analysis is not limited to a certain cell culture plate or culturing method, but can broadly be applied to fit various research needs.

**Figure 8:**
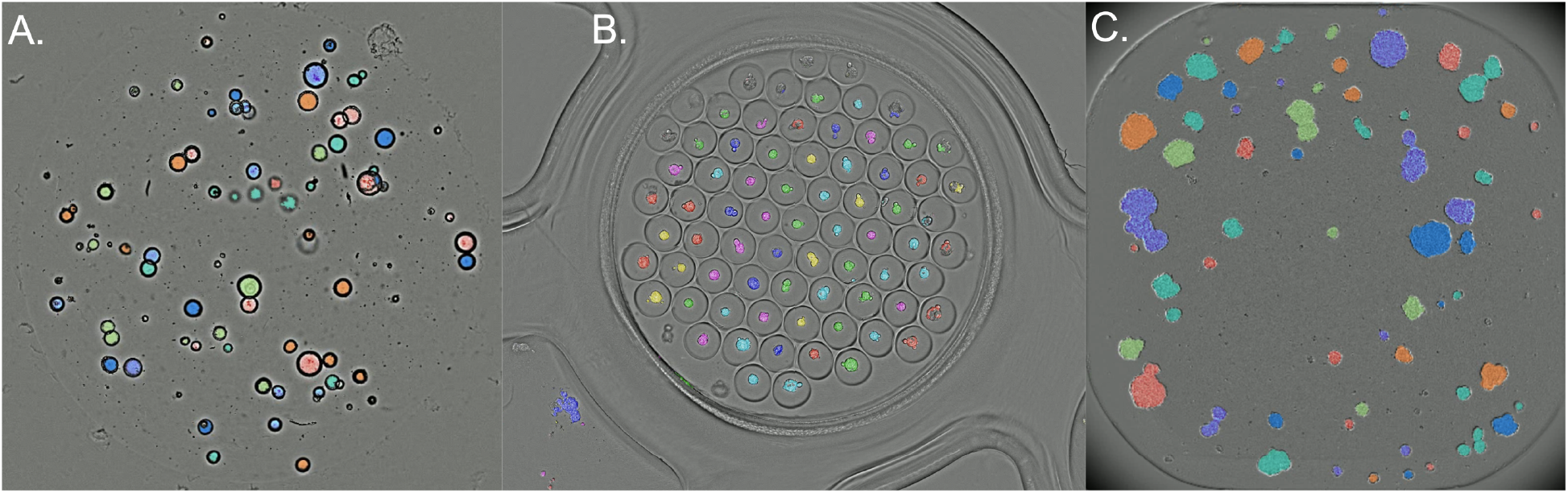
Examples of the broad applicability of OrBITS. **(A)** Cystic pancreatic cancer organoids (PDAC_060) grown in an extracellular matrix dome. Kinetic growth monitoring is shown in supplemental video 5. **(B)** Cancer cell line spheroids grown in a Gri3D^®^-96 well plate. Kinetic growth monitoring is shown in supplemental video 6. **(C)** NCI-H460 lung cancer cells grown as multispheroids and imaged with a 4x objective, phase-contrast using an IncuCyte S3 (Sartorius). Kinetic growth monitoring following treatment with cisplatin is shown in supplemental video 7. Colors indicate if connected objects are segmented individually (different colors), or as one object (same color).

Lastly, for broad applicability of the OrBITS analysis software, it was also crucial to demonstrate that it could be used on different image capturing systems. Therefore, the neural network was trained on images obtained with the Spark Cyto 600 system (Tecan), we also assessed its ability for masking with images acquired by the IncuCyte S3 (Sartorius). NCI-H460 lung cancer cells were grown as multispheroids and imaged with a 4x objective in phase-contrast. OrBITS was once again able to accurately mask the multispheroids and provide kinetic growth monitoring (Fig. 8C, Supplemental Video 7), thus indicating that the OrBITS analysis software is not limited to a single image capturing instrument.

Overall, we showed that the OrBITS software has broad applicability, in terms of various cell and organoid types, imaging systems and culturing methods, ranging from conventional to state-of-the art.

## 4. Discussion

In this study we validated a new brightfield imaging-based method for ex vivo drug screening on patient-derived organoids. The Spark Cyto 600 system used here is a state-of-the art multimode plate reader with BF and fluorescence imaging capabilities. With full environmental controls (e.g. O_2,_ CO_2_, temperature) and an injector, we automated the system to plate single cells or full-grown organoids, perform kinetic measurements, and dispense reagents (e.g. CellTiterGlo-3D) for endpoint analysis. This greatly improves the organoid culture by reducing variability involved with the cumbersome manual cultures, particularly in 384-well plates. While this all-in-one system ideal for the clinical implementation of drug sensitivity screens on patient-derived tumor organoids, the current limitation is the Spark Cyto Image Analyzer software, which is limited to 2D analysis. Therefore, we developed a deep learning image analysis solution OrBITS for 3D organoid/spheroid segmentation based on BF images.

Our system identifies organoids from BF images, thus removing the need for nuclear or cytoplasmatic fluorescent labeling, which can affect biological process and confound therapy responses. As demonstrated, there are several benefits to using the BF channel to detect and measure organoids over transient or stably expressed fluorescent markers. These include: (i) the ability to detect organoids in cases where high cell death reduces detectability via nuclear markers; (ii) increased reproducibility since variance related to dye concentrations, stability (e.g. photobleaching), LED intensity, and exposure time is removed; (iii) the ability to detect cystic organoids, which can be problematic since Hoechst only stains the outer edge of cystic organoids; (iv) optimal size and growth metrics, since it is not guaranteed that all parts of the organoid object will contain viable nucleated cells as organoids grow, and therefore, reliance of fluorescent markers therefore must be limited; (v) the ability to distinguish valid organoids from artifacts such as cellular debris which may produce false positives or bubbles which may occlude cells; (vi) and lastly, the reduction of any interfering effects of repeated fluorescence imaging on growth and metabolism due to the release of reactive oxygen species by photoexcited fluorophores and other side-effects related to phototoxicity (20).

Without the need to consider cytotoxic interference, imaging can be performed at higher frequency time intervals. This opens up new possibilities for studies into how drugs affect organoid growth and migration over time, which is currently not possible with simple endpoint assays. A key measurement parameter of drug screening studies is cell viability. This is commonly done using CellTiter-Glo 3D which, as an endpoint assay, is highly constrained. Alternatives to CellTiter-Glo 3D for measurement of cell death and viability are especially welcomed, since several therapies are known to affect the primary parameter of measurement for CellTiter-Glo 3D (ATP) via modulating intracellular ATP levels or releasing extracellular ATP following (immunogenic) cell death (21). Furthermore, this approach does not distinguish cytotoxic from cytostatic responses, which could further improve translatability of drug responses to the clinic and requires further investigation.

Using our method, we demonstrated that organoids can be grown from either single-cells or full-grown organoids with equal efficiency in 384-well plates. Gao et. al recently demonstrated that organoids starting from single-cells demonstrated similar sensitivity to cytotoxic drugs to matched full grown organoids (22). However, we recommend plating full grown organoids or at least delay treatment until multicellular organoids have been grown from the single cells. By using live-cell imaging, our method can greatly reduce turn-around time for *ex vivo* drug screenings as OrBITS can accurately monitoring organoid growth, health and therapy response with a limited amount of starting material on a single organoid level (13). In addition, kinetic imaging allows for early detection of therapy response and the use of growth rate metrics has been shown to result in higher reproducibility via uncoupling the effect of cell proliferation on drug sensitivity (14, 23). In contrast to CellTiter-Glo 3D, growth rates can be calculated for each well individually to reduce variations caused by seeding density. The use of a positive and negative control and the normalized drug response, as developed by Gupta and colleagues, resulted in the best discrimination in gemcitabine drug response between two PDAC patients with varying clinical outcomes following neoadjuvant gemcitabine-paclitaxel treatment. A larger study is currently ongoing in our group to support our hypothesis that the combination of live-cell imaging, OrBITS image analysis and the normalized drug response metric can improve stratification of patients into responders and non-responders.

Here, we demonstrated organoid identification, growth, and death using BF images, but we have only begun to tap into the potential of BF image analysis. In principle, other morphological changes with clinical and research implications are currently being pursued for organoids, 3D spheroid cultures, and 2D monolayers. We also demonstrated the flexibility of the OrBITS platform to differing culture methods, ranging from conventional (e.g. well-plates, matrix domes) to state-of-the-art (Gri3D^®^-96), and imaging systems (SparkCyto 600 and IncuCyte S3). Furthermore, we demonstrate the analysis capability on various cell and organoids lines apart from those used for developing the BF imaging software. In the process, we confirmed the challenges related to establishing lung cancer organoids as described by Dijkstra et al. (18). A more minimal medium compared to the one used in this study has recently been described which should limit the growth of normal lung organoids (24), although we observed no sustained organoid growth tested in a small set of samples. The use of Napsin A, p63 or p40 (ΔNp63) and TTF-1 allows for identifying cultures overgrown by non-malignant lung cells. As such, the patient-derived organoids used in our method development were non-malignant organoids displaying either a solid (derived from tumor tissue) or cystic (derived from healthy tissue) growth pattern. Apart from the clear difference in morphology, the non-malignant organoids derived from tumor tissue (NSCLC_051T) were more sensitive to cisplatin compared to the organoids derived from healthy tissue from the same patient (NSCLC_051N). This further indicates that these cells have different characteristics which could influence the outcome of a drug sensitivity screen if present in tumor organoid cultures. Even so, our method is also be applicable in tumor organoids as shown for PDAC organoids (18, 24).

## 5. Conclusions

We demonstrated that our in-house developed OrBITS image analysis software can monitor PDOs with high accuracy ranging from single cells to full-grown organoids based on BF imaging. Inclusion of a fluorescent cell death marker allowed for real-time monitoring of intra-well normalized organoid death. When validated against the current gold standard CellTiter-Glo 3D assay, the OrBITS imaging analysis was robust in determining cell viability, IC50-values. Furthermore, we demonstrated that OrBITS could distinguish cytostatic from cytotoxic responses most accurately by the inclusion of a fluorescent cell death marker and the use of the normalised drug response metric to provide more in-depth insight into anticancer drug mechanisms. Combined with providing high-throughput, kinetic monitoring of PDOs, OrBITS is a significant advance from the current state-of-the-art analysis technique made available for mid-end live-cell imaging instruments, which are often simple, endpoint analysis, or provide no mechanistic insights. Taken together, OrBITS, as a scalable, high-throughput technology, would facilitate the use of PDOs for drug development, therapy screening, and guided clinical decisions for personalized medicine. The developed platform also provides a launching point for further brightfield-based assay development to be used for fundamental research.

## List of Abbreviations

OrBITS: Organoid Brightfield Identification-based Therapy Screening
PDOs: Patient-derived organoids
BF: Brightfield
NSCLC: Non-small cell lung carcinoma
UZA: Antwerp University Hospital
RFU: Relative fluorescence units

## Declarations

### Ethics approval and consent to participate

Written informed consent were obtained from all patients, and the study was approved by the UZA Ethical Committee (ref. 17/30/339, 14/47/480).

### Consent for publication

All co-authors have read and approved of its submission to this journal and no individual person’s data is used.

### Availability of data and materials

The datasets used and/or analyzed during the current study are available from the corresponding author on reasonable request.

### Authors contribution

C.D, E.C.DLH. and A.L. conceived the idea and wrote the manuscript. C.D., M.L.C. performed the in vitro experiments. E.C.DLH. developed the software and analyzed the data. P.V.S., J.M.H, P.L, S.K.Y supplied the patient derived materials. F.L, P.P, A.B, E.S., A.L. substantially revised the manuscript.

## Acknowledgements

We would like to thank Mr. Willy Floren for funding the D300e and Tecan for gifting the Spark Cyto as part of a competition.

