## Supplementary Information, Tables and Figures for "OrBITS: Label-free and time-lapse monitoring of patient derived organoids for advanced drug screening"

### 1. Image analysis and training methodology

#### 1.1 Hoechst Organoid Detection

In order to compare object detection results, Hoechst fluorescence intensity images were segmented in a 3-step process: denoise, threshold, and label connected objects. These were used as an accepted truth for comparing count, mean area, and total area measurements.

To achieve ground truth from raw Hoechst fluorescent images, the images were first normalized from UINT8 to float 1.0 and median filtered with a 3x3 kernel. A gaussian blur was taken and subtracted from the image to remove low frequency signal. Images were binarized using a threshold value of 0.15. Individual organoids were indexed using connected component analysis. Objects with areas under 90 pixels$^2$ and above 7000 pixels$^2$ were removed. Holes with a size of less than 5000 pixels$^2$ were filled so that cystic organoids could be properly detected and measured.

#### 1.2 Using Fluorescence as Ground Truth

In order to use the Hoechst fluorescence intensity images as ground truth, we preprocessed the intensity values to be used against the predictions as a loss function similar to the above methods. Saturation clipping was used rather than thresholding, which provided the benefit of maintaining soft probabilities, thus minimizing the impact of poor threshold choice. This also provides better gradients for learning by back propagation. Soft probabilities are especially important as organoids increase in size while the total amount of Hoechst does not increase. Therefore, the dye distributed within the organoid results in a weaker signal over the course of growth.

#### 1.3 Model Architecture

A fully convolutional network was used to take single grayscale channel images as input of the BF channel and return sematic segmentation maps as output. These output maps were the same size in height and width as the input image and represent a predictive response for each pixel and class in the input image.

#### 1.3.1 Detection vs Classification Architecture

A key change in our model architecture was that it functions as a detector rather than a classifier. This allowed us to bypass the SoftMax function which was used as the final step of the original UNET. The benefits of this decision were two-fold: first, it does not require the learning and evaluation of background as a class, making it possible for previously unseen objects (e.g. cellular debris, glare, shadows) to be ignored without dedicated training, and second, it allows learning and production of multiple, simultaneous outputs in which a single pixel may be positive in multiple classes, (e.g. detection of location, detection of key behaviors such as death and mitosis, detection of subcomponents such as filopodia, nuclei, and cystic).

#### 1.3.2 Implementation Resources

Our neural network was implemented in python 3.8 using PyTorch 1.8.0. GPU acceleration was enabled for training using CUDA 11, though full completion of training was possible under 20 minutes, even without acceleration, using an Intel CORE i7 Processor. Training required at most 1000 cycles to reach a convergence at mean DICE loss of .15 (16).

### 2. Challenges in establishing lung cancer organoids

We faced the same challenges in generating non-small cell lung cancer (NSCLC) organoids as described by Dijkstra and colleagues [12]. Organoids derived from tumor resection fragments had a predominant solid growth pattern, while organoids derived from distant healthy lung tissue displayed a cystic growth pattern (Fig S1). Immunohistochemistry (IHC) characterization showed a polarized staining pattern for antibody p40 (ΔNp63), an isoform of p63 which is seen in the stem-like cell population of basal cells in the respiratory epithelium and which expression is lost in more differentiated cells [13]. P40 is considered a more selective marker for distinguishing lung squamous cell carcinoma (+) from adenocarcinoma (-) compared to p63. These findings are consistent with the findings of Dijkstra et al., showing polarized expression of p63 in tumor-derived organoids with a normal copy number profile. Together with the absence of Napsin A in the adenocarcinoma samples (NSCLC_006T, NSCLC_013T, NSCLC_046T) and TTF-1 positivity in the squamous cell carcinoma sample (NSCLC_051T), this indicates that these are non-malignant cells and most likely represent metaplastic squamous epithelial cells [12,14] (Fig. S1). For NSCLC_006T and NSCLC_013T this was genetically confirmed since the organoids lack the parental KRAS and EGFR mutations, respectively (Table 1).

### 3. Supplemental Tables

| Table S1: Cytometry Metrics | | | |
|---|---|---|---|
| **Detection** | | | |
| Object Counts | **N** | The number of nonconnected objects identified as organoids. [1] | - |
| **Size and Growth** | | | |
| Mean Mask Area | $\boldsymbol{A_\mu}$ | The average area of each organoid in the well. [1] | $\frac{1}{N}\sum_{i}^{N} P_i$ |
| Total Mask Area | $\boldsymbol{A_\Sigma}$ | The sum area of all organoids in the well<br>This metric is equivalent to the Mean Mask Area * Object count. | $\sum_{i}^{N} P_i$ |
| **Signal Intensity** | | | |
| Mean Intensity | $\boldsymbol{I_\mu}$ | The average of the [average fluorescence intensity in each organoid] for all organoids in the well. [1] | $\frac{1}{N}\sum_{i}^{N}\frac{\sum_{j}^{P_i} I_{ij}}{P_i}$ |
| Sum Intensity | $\boldsymbol{I_\Sigma}$ | The sum of the average fluorescence intensity for all organoids in the well. | $\sum_{i}^{N}\frac{\sum_{j}^{P_i} I_{ij}}{P_i}$ |
| Total Mask Area – Total Intensity | - | The difference between the total mask area in the well and the total intensity. | $\boldsymbol{A_\Sigma} - \boldsymbol{I_\Sigma}$ |
| **Cell Viability** | | | |
| Mean Signal Positivity | $\boldsymbol{\Phi_\mu}$ | The average organoid positivity for all organoids in the well. Where positivity is a binary if the mean signal organoid is above a given threshold. (τ) [1,2] | $\phi = \frac{1}{P_i}\sum_{j}^{P} I_{ij} > \tau$<br>$\frac{1}{N}\sum_{i}^{N}\phi$ |
| Total Signal Area | $\boldsymbol{I_{A\Sigma}}$ | The sum of the number fluorescence values that are above a threshold | $\sum_{i}^{N}\sum_{j}^{P_i}(\mathbf{I_{ij}} > \tau)$ |
| Total Signal Area / Total Mask Area | - | The total signal area divided by the total mask area.[2] | $\boldsymbol{I_{A\Sigma}} / \boldsymbol{A_\Sigma}$ |
| Total Mask Area – Total Signal Area | - | The difference between the total mask area in the well and the total signal area.[2] | $\boldsymbol{A_\Sigma} - \boldsymbol{I_{A\Sigma}}$ |
| 1 This metric requires individual object detection.<br>2 This metric requires setting an adjustable threshold value. | | | |

| Table S2: Correlation Results Summary | | | | | | | |
|---|---|---|---|---|---|---|---|
| | | **Untreated - Variable Seeding** | | | **Cisplatin Treated** | | |
| | | Slope | Bias | $R^2$ | Slope | Bias | $R^2$ |
| **Detection** | | | | | | | |
| Object Counts | **N** | 0.89 | 4.76 | .98 | 0.67 | 5.74 | .79 |
| **Measuring size and growth** | | | | | | | |
| Mean Mask Area | $\boldsymbol{A_{\mu}}$ | 1.01 | 45.1 | .93 | 0.74 | 34.7 | .66 |
| Total Mask Area | $\boldsymbol{A_{\Sigma}}$ | 1.04 | 4211 | .95 | 0.76 | 577 | .78 |
| **Measuring Cytotox Signal Intensity** | | | | | | | |
| Mean Cytotox Area | $\boldsymbol{I_{\mu}}$ | - | - | - | 0.92 | 0.04 | .92 |
| Mean Cytotox Signal | $\boldsymbol{I_{\Sigma}}$ | - | - | - | 0.85 | 0.02 | .94 |

| Table S3: Overview of compatible cancer cell lines | | | |
|---|---|---|---|
| Cell line | **Spheroids in ECM dome** | **Spheroids in 384-well** | **Comment** |
| A549 | Yes | Yes, static | |
| Calu-3 | Yes, slow | Yes, merge | Do not plate to dense, use 3-day old organoids |
| HCT-116 | Yes | Yes, static | |
| SW480 | No | Yes, merge into 1 large spheroid if plated to dense | Plate <500 cells, allow to grow for 2 days in the 384-well before starting treatment. |
| Panc-01 | Yes | Yes | |
| BxPC3 | Yes | Yes | |
| MiaPaca2 | No | Yes, merge | Plate <500 cells, allow to grow for 2 days in the 384-well before starting treatment. |
| NCI-H1975 | Yes, but 2D-outgrowth. Grow at least 1 passage in domes. | Yes, after >1 passage cultured in domes | |
| NCI-H82 | No | No | Suspension cells |
| HCT-15 | Yes | Yes | |
| NCI-H460 | Yes | Yes | |
| SCC22b | Yes | Yes | |
| SC263 | Yes | Yes | |
| SQD9 | Yes | Yes | |
| LICR-HN1 | Yes | Yes | Easily merge together so try plating <500 cells in 384-well ULA plate and grow for 2 days instead of in dome. |
| Static: spheroids are not migratory. Merge: Spheroids migrate towards each other and merge together. | | | |

### 4. Supplemental Figures

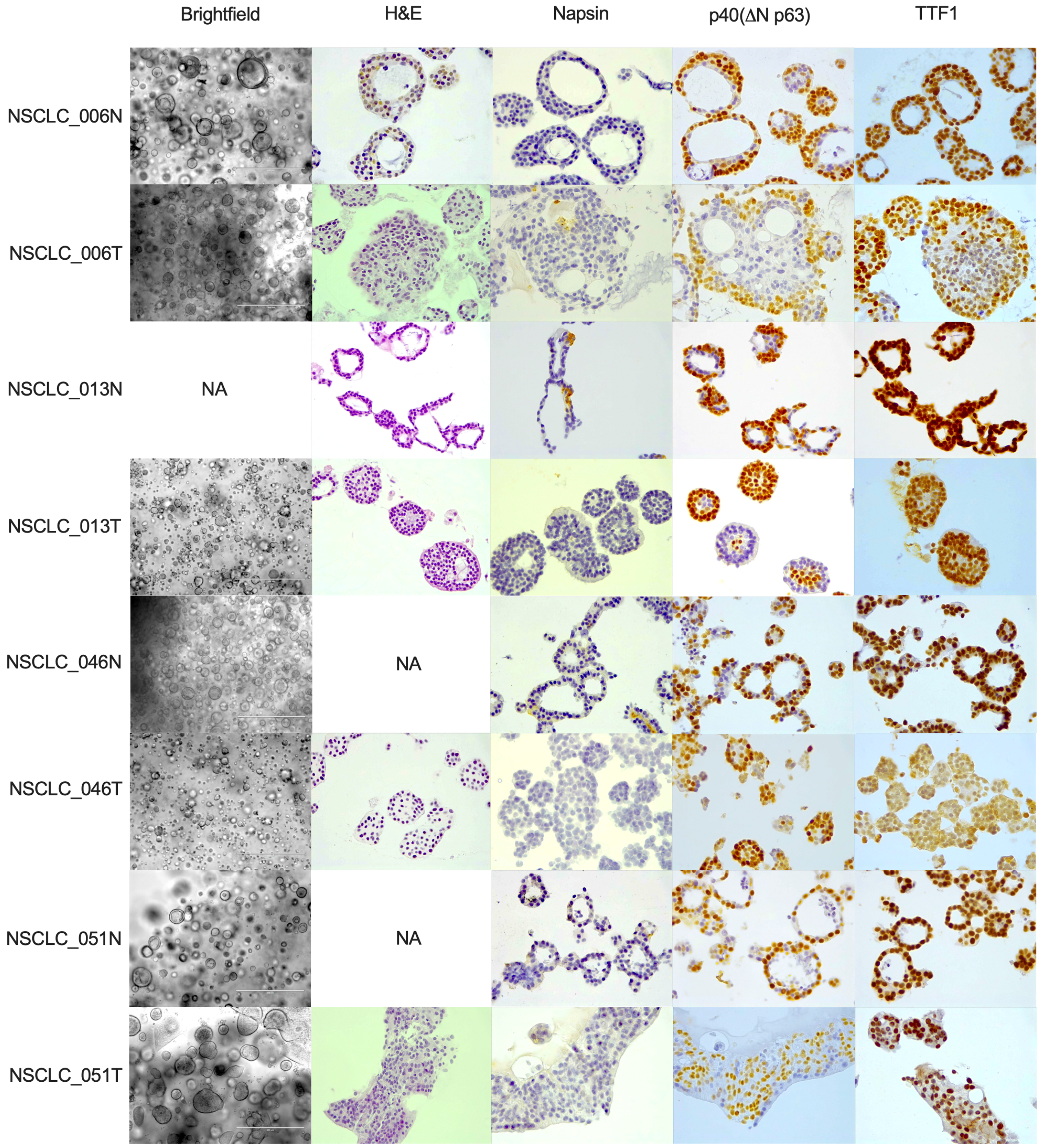

**Figure S1:** Overview of the patient-derived organoid (PDO) morphology and expression of lung (cancer) related markers by immunohistochemical staining.

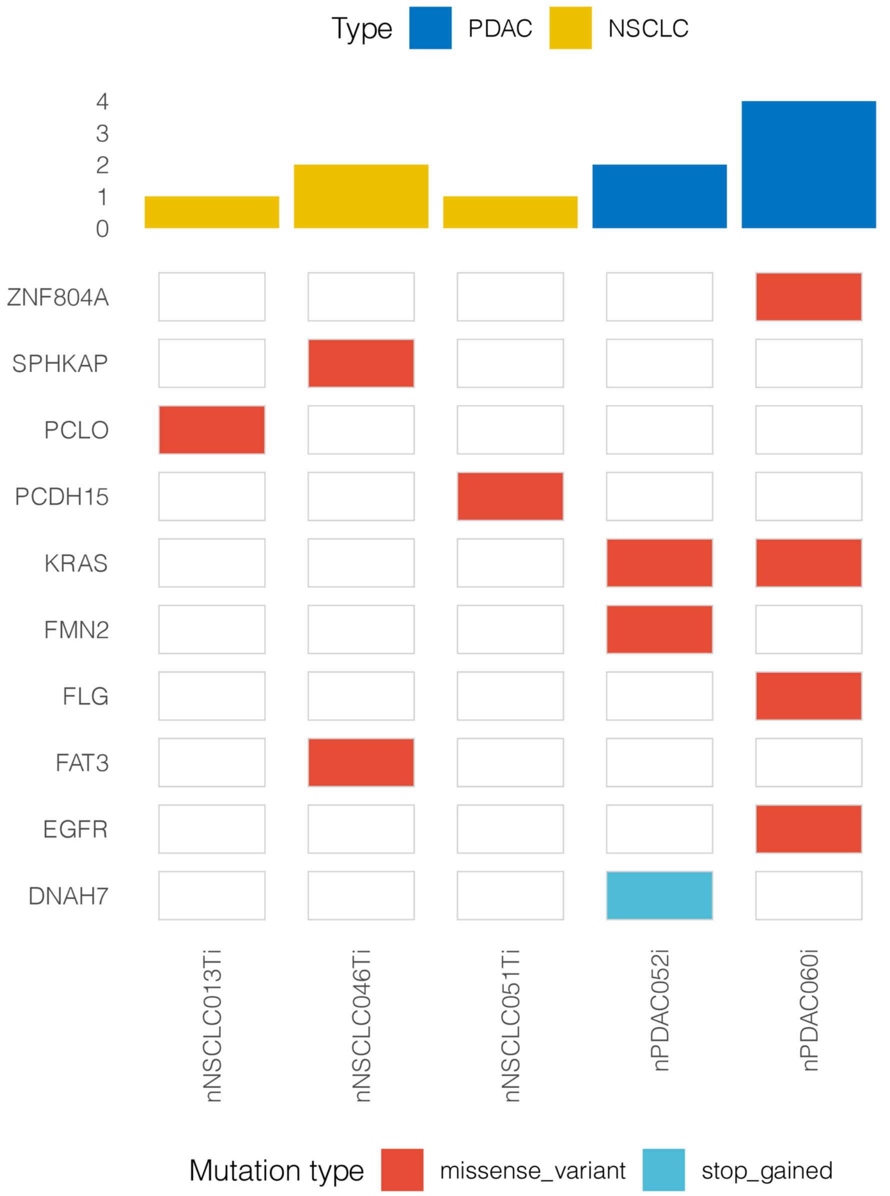

**Figure S2:** Oncoprint NSCLC and PDAC organoids. Only genes are shown in which mutations were detected. No mutations were detected in NSCLC_006N.

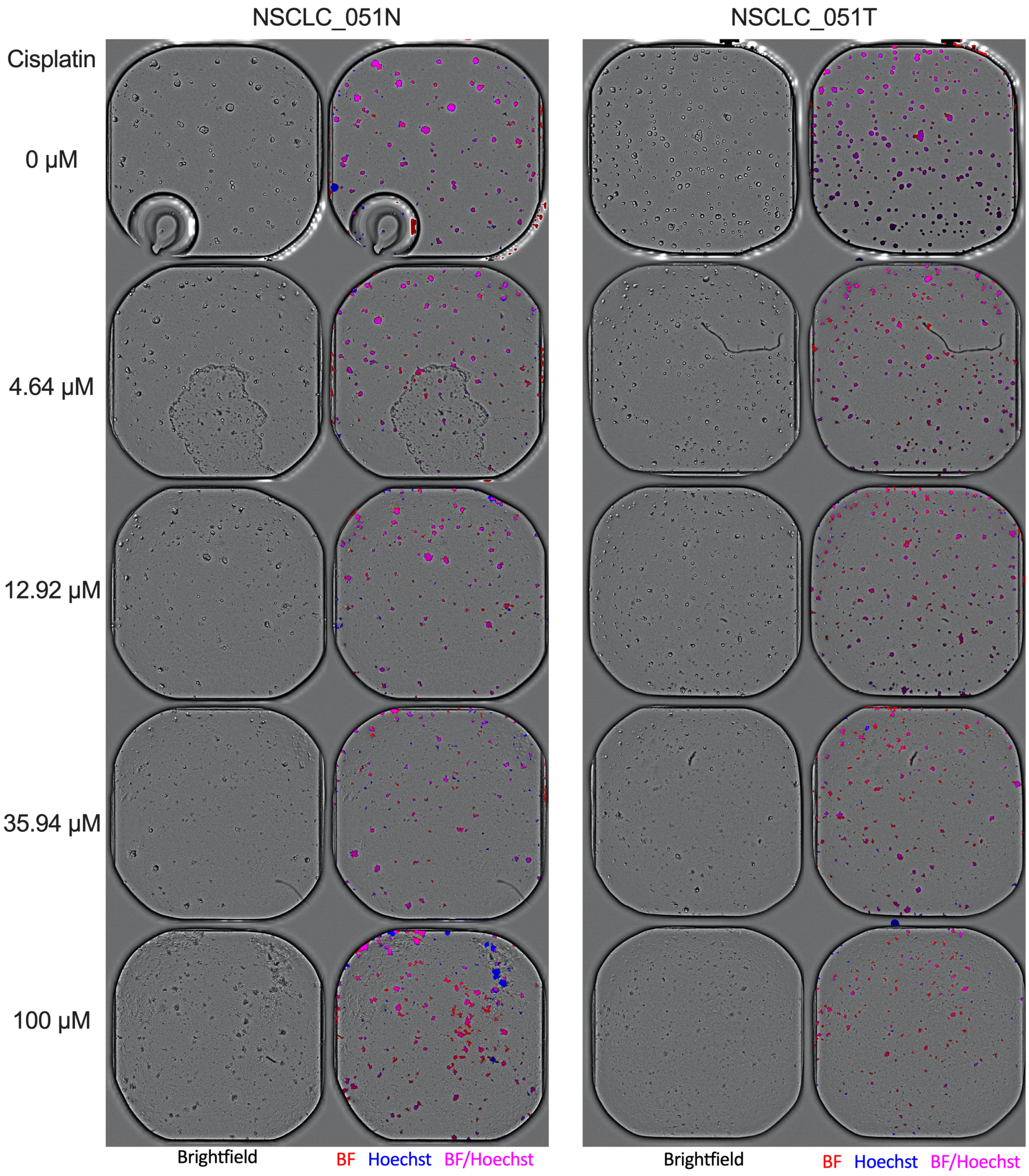

**Figure S3: Validation of OrBITS BF organoid tracking compared to Hoechst.** Representative images of NSCLC_051N and NSCLC_051T PDOs treated with different concentrations of the chemotherapeutic drug cisplatin shows that organoids with varying levels of cytotoxicity can be identified with both Brightfield- and Hoechst-based analysis. In addition, the figures show that common artefacts (e.g. air-bubbles, extracellular matrix, dust) were excluded from analysis.

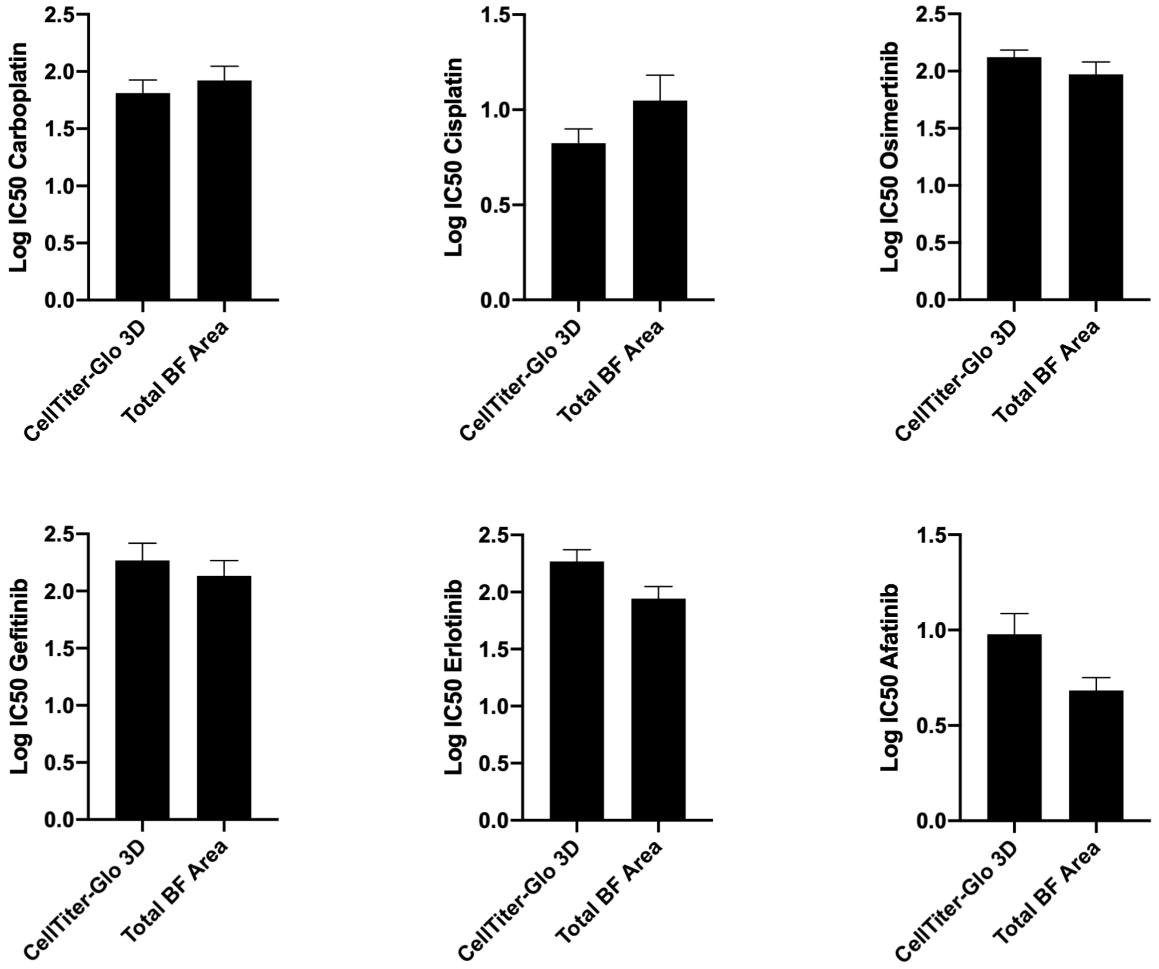

**Figure S4:** Log IC50 values for NSCLC_013T treated with a 7-point titration of cisplatin, carboplatin, erlotinib, gefitinib, osimertinib or afatinib normalized to vehicle (100%) and baseline (0%, 1 μM staurosporine).

**4. Supplemental Video Legends**

**Supplemental video 1:** Representative time-lapse video (4h interval) of whole 384-well brightfield (top left) and fluorescent images (top right) of 3-day old NSCLC_013T organoids seeded a 1000 organoid/wells. The multi-colored mask identifies organoids from BF imaging (bottom left). OrBITS also allows for tracking of the organoid movement over time (bottom right). Blue fluorescence = Hoechst, green fluorescence = cytotox green, coloured mask = BF-based masking.

**Supplemental video 2:** Representative time-lapse video (4h interval) of whole 384-well brightfield (top left) and fluorescent images (top right) of NSCLC_006N organoids seeded at a density of 500 single cells/wells. The multi-colored mask identifies organoids from BF imaging (bottom left). OrBITS also allows for tracking of the organoid movement over time (bottom right). Blue fluorescence = Hoechst, green fluorescence = cytotox green, coloured mask = BF-based masking.

**Supplemental video 3:** Representative time-lapse video (6h interval) of whole 384-well brightfield (left) and fluorescent images (middle) of full-grown PDOs and the corresponding analysis mask (right). Blue fluorescence = Hoechst, green fluorescence = cytotox green, red mask = BF, pink mask = BF/Hoechst overlap, white mask = cytotox green.

**Supplemental video 4:** Example of kinetic measurement (4h interval) of the growth of cancer cell lines spheroids or PDOs from brightfield images. Panel 1: PDOs from pancreatic cancer, panel 2: NCI-H1975, panel 3: HCT-15, panel 4: NCI-H460. Growth is shown as total BF mask area and cell death as total Green mask area based on cytotox green.

**Supplemental video 5:** Example of the brightfield-based masking of cystic pancreatic cancer organoids (PDAC_060) grown in extracellular matrix domes. Images were acquired every 2 hours using a 2x objective.

**Supplemental video 6:** Example of brightfield-based masking of spheroids grown in the Gri3D®-96 well plates. Images were acquired every 4 hours using a 4x objective.

**Supplemental video 7:** Example of phase-contrast images of NCI-H460 lung cancer spheroids treated with vehicle, 2.5µM, 6.3µM and 15.5µM cisplatin. Images were acquired every 4 hours using a 4x objective.
